# The hormone neuroparsin seems essential in Lepidoptera but not in domesticated silkworms

**DOI:** 10.1101/716746

**Authors:** Jan A. Veenstra

## Abstract

The primary sequence of the Arthropod neurohormone neuroparsin is so variable that so far no orthologs from moths and butterflies have been characterized, even though classical neurosecretory stains identify cells that are homologous to those producing this hormone in other insect species. Here Lepidopteran cDNAs showing limited sequence similarity to other insect neuroparsins are described. That these cDNAs do indeed code for authentic neuroparsins was confirmed by *in situ* hybridization in the wax moth, *Galleria mellonella*, which labeled the neuroparsin neuroendocrine cells. Although in virtually all genome assemblies from Lepidoptera a neuroparsin gene could be identified, the genome assembly from the silkworm, *Bombyx mori*, has a neuroparsin gene containing a 16 nucleotide deletion that renders this gene nonfunctional. Although only a small number of all silkworm strains carry this deletion, it suggests that the domestication of the silkworm has rendered the function of this neurohormone dispensable.

## 1. Introduction

In a 1928 publication Ernst Scharrer described what are now known as neuroendocrine cells producing vasopressin and oxytocin (Scharrer, 1928). At the time the concept of what was later called neurosecretion and neuroendocrinology, but what at the time he called internal secretion (“innere Sekretion”), was not readily accepted by vertebrate physiologists. However, insect physiologists had already demonstrated the existence of a “brain hormone” that activates the prothoracic gland (*e.g.* Kopeć, 1922) and this strongly stimulated research on neurosecretory cells in insects. One of the most popular staining methods uses paraldehyde fuchsin, which stains several cells in the insect *pars intercerebralis* that have axons projecting to the *corpora cardiaca*, where they release their products into the hemolymph. There are generally two different paraldehyde fuchsin cell types in the *pars intercerebralis*, which differ by their staining intensity. We now know that the less intensely staining cells produce insect insulins and the other neuroparsins (Goltzené et al., 1992).

The structure of neuroparsin was first identified from *Locusta migratoria* (Girardie et al, 1989). Nevertheless, relatively little is known about the physiological relevance of this hormone in insects, no doubt due in large part to its absence from *Drosophila melanogaster* (Veenstra, 2010), the insect neuropeptide model par excellence. In *Locusta migratoria* the hormone antagonizes juvenile hormone. It delays vitellogenesis and immunoneutralization of neuroparsin accelerates vitellogenesis and induces green pigmentation. These effects are similar to those of juvenile hormone (Girardie et al., 1987). Once its cDNA was cloned, neuroparsin was found to be expressed also outside the central nervous system (Lagueux et al., 1992) and the identification of several distinct cDNAs showed that they were derived from a single gene that is alternatively spliced. The genome of *L. migratoria* revealed the neuroparsin gene to have seven coding exons that allow the production of five different neuroparsin transcripts (Veenstra, 2014). Work on another migratory locust, *Schistocerca gregaria*, revealed a very similar situation, although the genome for this species has not yet been sequenced and so far only four different neuroparsin transcripts have been identified (Girardie et al., 1998; Janssen et al., 2001; Claeys et al., 2003, 2005, 2006a,b). Interestingly, expression of the different transcripts is significantly different between the solitary and gregarious phases in the latter species (Claeys et al., 2006a).

The extensive alternative splicing of neuroparsin transcripts seems to be limited to migratory locusts, as to my knowledge it has not been found in other Arthropods. Some species, notably various Decapods (Veenstra, 2016) and blood sucking bugs like *Rhodnius prolixus* (KQ034340.1) and *Cimex lectularis* (Predel et al., 2018), however have multiple neuroparsin genes. In this respect neuroparsin resembles insulin, which in many species also has multiple genes. This is not the only similarity between neuroparsin and insulin. Thus, both neuroparsin and insulin promote neurite outgrowth in locust brain (Vanhems et al., 1990). Furthermore, identification of ovarian ecdysteroid hormone from the mosquito *Aedes aegypti* revealed it to be a neuroparsin ortholog (Brown et al., 1998) and the subsequent identification of a venus kinase as its receptor (Vogel et al., 2015) showed that insulin and neuroparsin use similar receptors in insects. In mosquitoes both hormones stimulate the ovary to produce ecdysteroids which in turn trigger vitellogenin synthesis by the fat body (Dhara et al., 2003). Thus these two hormones seem somewhat complimentary. This is also suggested by venus kinase RNAi injections in *S. gregaria*, where both neuroparsin and insulin transcripts are upregulated. Downregulation of the venus kinase receptor by RNAi leads to a delay in vitellogenin synthesis and strongly inhibits production of ecdysone and several of the enzymes needed for its synthesis. Nevertheless, overall reproductive success appeared to be only slightly impacted in *Schistocerca* (Lenaerts et al., 2017).

The brains of Lepidoptera also typically contains two cell types that are stained by paraldehyde fuchsin (Panov and Kind, 1963) and Lepidopteran insulins have been extensively studied in the silkworm (for review see: Mizoguchi and Okamoto, 2013). However, in spite of much research on neuropeptides in the silkworm, and the presence of two venus tyrosine kinase receptors in its genome (Vanderstraete et al., 2013), their putative ligands have so far escaped identification. I here show that neuroparsin seems ubiquitously present in Lepidoptera but that in some domesticated silkworm strains the gene has mutated in such fashion that it can no longer be released as a hormone into the hemolymph.

## 2. Materials and Methods

### 2.1. Bioinformatics

After the initial identification of the *Galleria* and *Bombyx* neuroparsin genes 63 lepidopteran genome assemblies available at NCBI (https://www.ncbi.nlm.nih.gov/genome/?term=lepidoptera) were analyzed for the presence of orthologous genes. Predicted neuroparsin precursors were deduced and the size of the intron between the first and second coding exons was determined. In most cases the three coding exons were found on the same contig, but in several genome assemblies the contigs are very small and in those cases the the coding exons were connected based on homology to produce a neuroparsin precursor. In four assemblies no neuroparsin gene was found, in two of these species, *Heliconius hecale* and *Heliothis virescens*, the genes were assembled from individual reads recovered from genomic SRAs, SRR7162651 and SRR5463746 respectively, using the sratoolkit (www.ncbi.nlm.nih.gov/sra/docs/toolkitsoft/) and Trinity (Grabherr et al., 2011) using a method described previously (Veenstra, 2019).

### 2.2 In situ *hybridization*

The *in situ* hybridization protocol is based on a protocol provided by Dušan Žitňan for *Drosophila* as described *e.g.* here (Kim et al., 2006). Dissections were done in 0.9 % NaCl and tissues were fixed in phosphate buffered 4% paraformaldehyde for 2 hrs at room temperture in Eppendorf tubes. Tissues were subsequently washed thrice for 30 min in PBS with 0.2 % Tween 20 (PBST) and once for 30 min in 70% ethanol. After replacing the 70% ethanol the tissues were stored at −20 °C for three to five days. Tissues were next washed thrice for 25 min in PBST and then incubated with Proteinase K (50 µg/ml) in PBST for 30 min. The proteinase K was stopped by washing the tissue for 30 min in glycin (2 mg/ml) in PBST, followed by two washes of 20 min in PBST. Next tissues were fixed a second time in phosphate buffered 4% paraformaldehyde for 1 hr followed by two washes in PBST for 20 min each. This was followed by incubations for 20 min in 1:1 mixture of hybridization solution (HS: 50 µg/ml heparin, 100 µg/ml salmon testes DNA, 750 mM NaCl, 75 mM sodium citrate, in deionized RNase free water, pH 7.0, 50% deionized formamide, 0.1 % Tween 20) and PBST, followed by 25 min in HS, both at roomtemperature. HS was then replaced with fresh HS and the tissues were incubated for 3 hrs at 48 °C, after which HS was replaced with fresh HS containing 10% of digoxygenin hybridization probe which had previously been brought to 95 °C for either 3 min, in case of a previously used probe, or 45 min when a hybridization probe is used for the first time. Hybridization was carried out overnight at 48 °C. The following day hybridization probe was recovered and stored at −20 °C for future use. Tissues are washed in fresh HS four times, thrice for 2 to 2.5 hrs, and then overnight, all washes in HS were done at 48 °C. The next day tissues were brought to room temperature and washed in a 1:1 mixture of PBST and HS, follwed by three 20 min washes in PBST at room temperature. Next tissues were saturated with 1% BSA in PBST for 1 hr. The 1% BSA in PBST was then used to dilute sheep anti-digoxygenin Fab-fragments conjugated to alkaline phosphatase (1:1000, Sigma-Aldrich) in which the tissues were incubated overnight at room temperature. Next morning tissues were washed thrice 20 min with PBST, followed by three washes in freshly prepared alkaline phosphate buffer (APB, 100 mM Tris, 50 mM MgCl2, 100 mM NaCl, pH 9.5 and 0.1% Tween 20). During the last wash, the tissues were transferred to small glass dissection dishes and then the location of the digoxygenin probe was visualized by replacing the wash solution with APB containing 20 µl/ml of a commercial NBT/BCIP stock solution (Sigma-Aldrich, 18.75 mg/ml nitro blue tetrazolium chloride and 9.4 mg/ml 5-bromo-4-chloro-3-indolyl-phosphate, toluidine-salt in 67% DMSO). Color development was followed under a dissecting microscope and took about five minutes. Once color development was judged satisfactory, the alkaline phosphatase was stopped by changing the staining solution with 100 mM phosophate buffer, pH 7.0. This was followed by four 15 min washes in increasing concentrations of glycerol (20 %, 40 %, 60 % and 80 %) and finally tissues were mounted between a slide and a coverslip in 80 % glycerol. All incubations at room temperature were performed on gently rotating orbital shaker.

### 2.3. Hybridization probes

RNA was extracted and reverse transcribed using random primers and M-MuLV reverse transcriptase (New England Biolabs) and a PCR product of about 300 nucleotides from a coding sequence was than amplified by PCR using Q5® High-Fidelity DNA polymerase (New England Biolabs). The resulting PCR products were gel purified, quantified and their identities as authentic neuroparsin cDNAs confirmed by automated Sanger sequencing. Sense and anti-sense digoxigenin labeled probes were then produced using Taq polymerase (New England Biolabs) and 40 cycles of PCR with either the sense or anti-sense primer in a Taq polymerase PCR mixture to which 0.067 mM of Digoxigenin-5-aminoallyl-dUTP (Jena Bioscience GmbH, Jena, Germany) had been added and in which the concentration of dTTP had been lowered to 0.133 mM. The final PCR product was used directly in the *in situ* hybridization procedure.

### 2.4. Primer sequences used

*Galeria* neuroparsin

forward primer 5′-CGAGTGTCCAGTATGCATCG-3′

reverse primer 5′-TTGGGATTAAGGAGCCATTG-3′

*Bombyx* neuroparsin

forward primer 5′-GAGAGACGAACCGGAAATCA-3′

reverse primer 5′-CCATGCCTTGACAGATACCC-3′

### 2.5. Insects

*Galleria mellonella* larvae were purchased from St. Laurent (La Chapelle Saint Laurent, France), while *Bombyx mori* pupae were a generous gift from Michal Žurovec from the University of Southern Bohemia.

## 3. Results

Extensive use of the BLAST program led to the identification of a neuroparsin precursor candidate that had very limited sequence homology with its orthologs from other species (Fig. 1), but that within the Lepidoptera is relatively well conserved (Fig. 2 and Table S1). It typically has a signal peptide and analysis of a limited number of RNAseq SRAs from *B. mori* (Table S2) suggested it to be relatively specific for the brain.

**Fig. 1.**
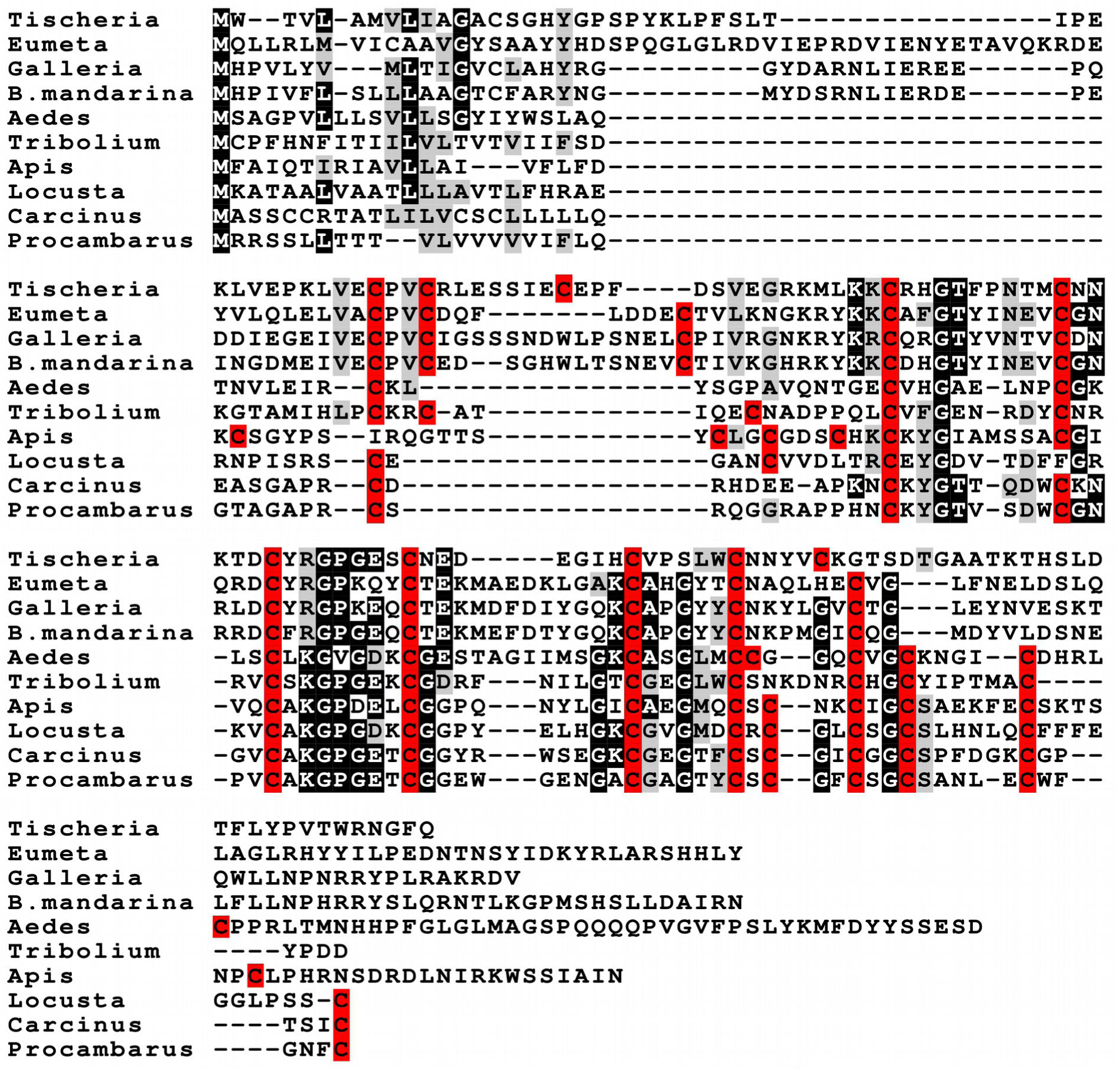
Alignment of selected Arthropod neuroparsin precursor sequences. Sequences are from *Tischeria quercitella* (GENO01134828.1), *Eumeta japonica* (BGZK01000877.1; join[241322..241488,242249..242387,243576..243797]), *Galleria mellonella* (XM_026905025), *Bombyx mandarina* (XM_028184972.1), *Aedes aegypti* (U69542.1), *Tribolium castaneum* (XM_015980588), *Apis mellifera* (XM_026440542), *Locusta migratoria* (Y17710.1), *Procambarus clarkii* (GARH01002923.1) and *Carcinus maenas* (GFYV01020843.1). Note the very limited sequence similarity of the Lepidopteran neuroparsins with those from other Arthropods. Primary sequence similarity is so poor that Clustal Omega no longer aligns the cysteine residues, which are highlighted in red; other conserved amino acid residues are highlighted in black.

**Fig. 2.**
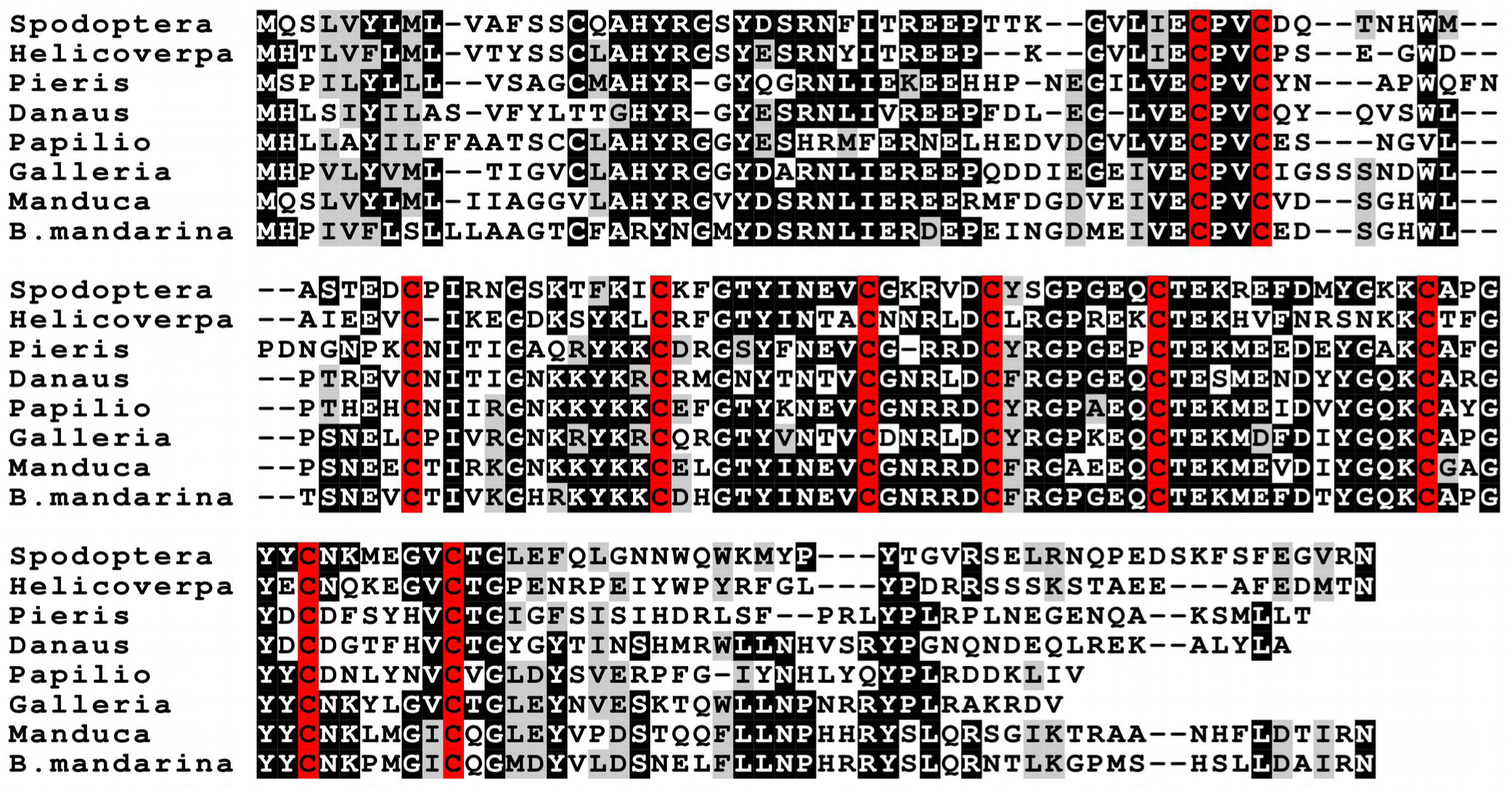
Alignment of selected Lepidoptera neuroparsin precursor sequences. Sequences are from *Spodoptera litura* (GBBY01013502.1), *Helicoverpa armigera* (GFWI01343101.1), *Pieris rapae* (GGJY01013372.1), *Danaus plexippus* (OWR50203.1), *Papilio machaon* (XP_014372220.1), *Galleria mellonella* (XM_026905025), *Manduca sexta* (GETI01101303.1) and *Bombyx mandarina* (XM_028184972.1). For more sequences see Table S1. Cysteine residues are highlighted in red and other conserved amino acid residues in black.

Several of the Lepidopteran genome assemblies are little more than first drafts and it is therefore not surprising that I was unable to find evidence for a neuroparsin gene in *Operophtera brumata*. On the other hand there is abundant genomic and transcriptome data for *Plutella xylostella* (see *e.g.* http://dbm.dna.affrc.go.jp/px/). This species is one of two studied here that does not belong to the Apoditrysia taxon. The other one is *Eumeta japonica*, and its neuroparsin precursor sequence is very different from that of the Apoditrysia (Fig. 1). Hence it seems plausible that the *Plutella* neuroparsin sequence was not recognized because it is too different from that of the other Lepidoptera.

To confirm or invalidate the hypothesis that these sequences might represent neuroparsin orthologs, whole mount *in situ* hybridization of brains was performed in *Galleria mellonella* larvae as well as adults. RNA extraction of *Galleria* heads and is subsquent reverse transcription allowed the production of a neuroparsin PCR product the identity of which was confirmed by Sanger sequencing (Figs. S1,S2). In both larvae and adults *in situ* hybridization revealed two bilateral groups of four large cells in the *pars intercerebralis* (Fig. 3) in the location where neuroparsin expressing cells were expected based on paraldehyde fuchsin staining reports (Panov and Kind, 1963). RNA extraction of a single *Bombyx* brain and is subsquent reverse transcription similarly allowed the production of a neuroparsin PCR product that was sequenced and found to have the 16 base pair solution (Figs. S3,S4). Although *in situ* hybridization procedures were identical to those performed on *Galleria*, which were run in parallel, no *in situ* hybridization signal was obtained in *Bombyx* adult brains. This could be due to low expression levels of neuroparsin in *B. mori*, as suggested by the 0.00 TPM (transcripts per million) values for the two brain RNAseq transcripts present at silkbase (http://silkbase.ab.a.u-tokyo.ac.jp). Digoxigenin probes generated with the sense probes did not yield any signal in either species.

**Fig. 3.**
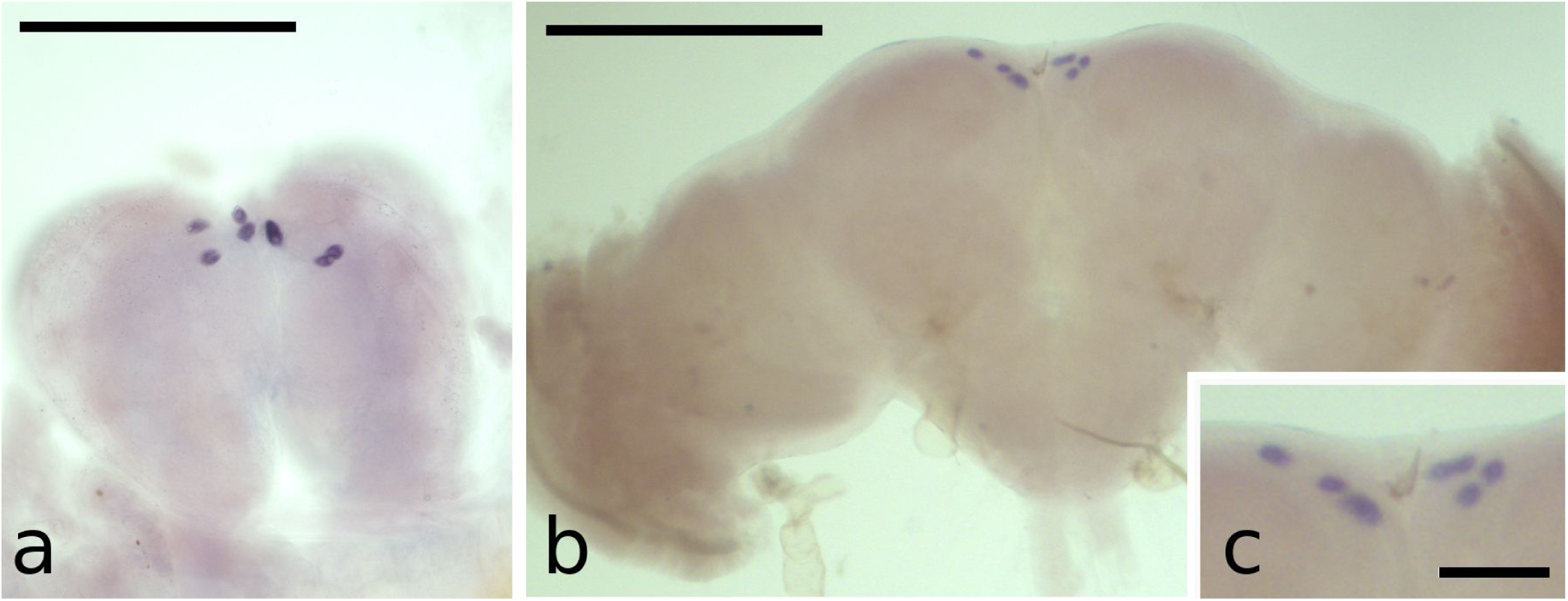
*In situ* hybridization of neuroparsin mRNA in the brain of *Galleria mellonella* larva **(a)**, and adult **(b)** with an amplification of the adult *pars intercerebralis* in **(c)**. Note that these are whole mounts and consequently it is difficult to get all cells in focus in a single picture. Scale bars are 500 μm in **(a)** and **(b)** and 200 μm in **(c)**.

Intriguingly in the *Bombyx mori* genome assembly, the neuroparsin gene does not code for a signal peptide. However, a signal peptide is predicted from the orthologous *B. mandarina* gene (Fig. 4), the species from which *B. mori* was domesticated about 5,000 years ago (Sun et al., 2012; Chen et al., 2019). There are two significant differences between the two genes. The first one concerns the intron between the first and second coding exons. In the domesticated silkworm genome assembly it contains 9,493 but that of the wild silkworm has only 827 base pairs. The second and more important difference is a unique deletion of 16 base pairs from the first coding exon (Fig. 4). Analysis of *B. mori* brain RNAseq SRAs yielded a total of 4,358 spots and Trinity unambiguously confirmed the proposed mRNA. This mRNA contains the unique deletion, as was expected since these RNAseq SRAs are all obtained from the same strain as used for genome sequencing.

**Fig. 4.**
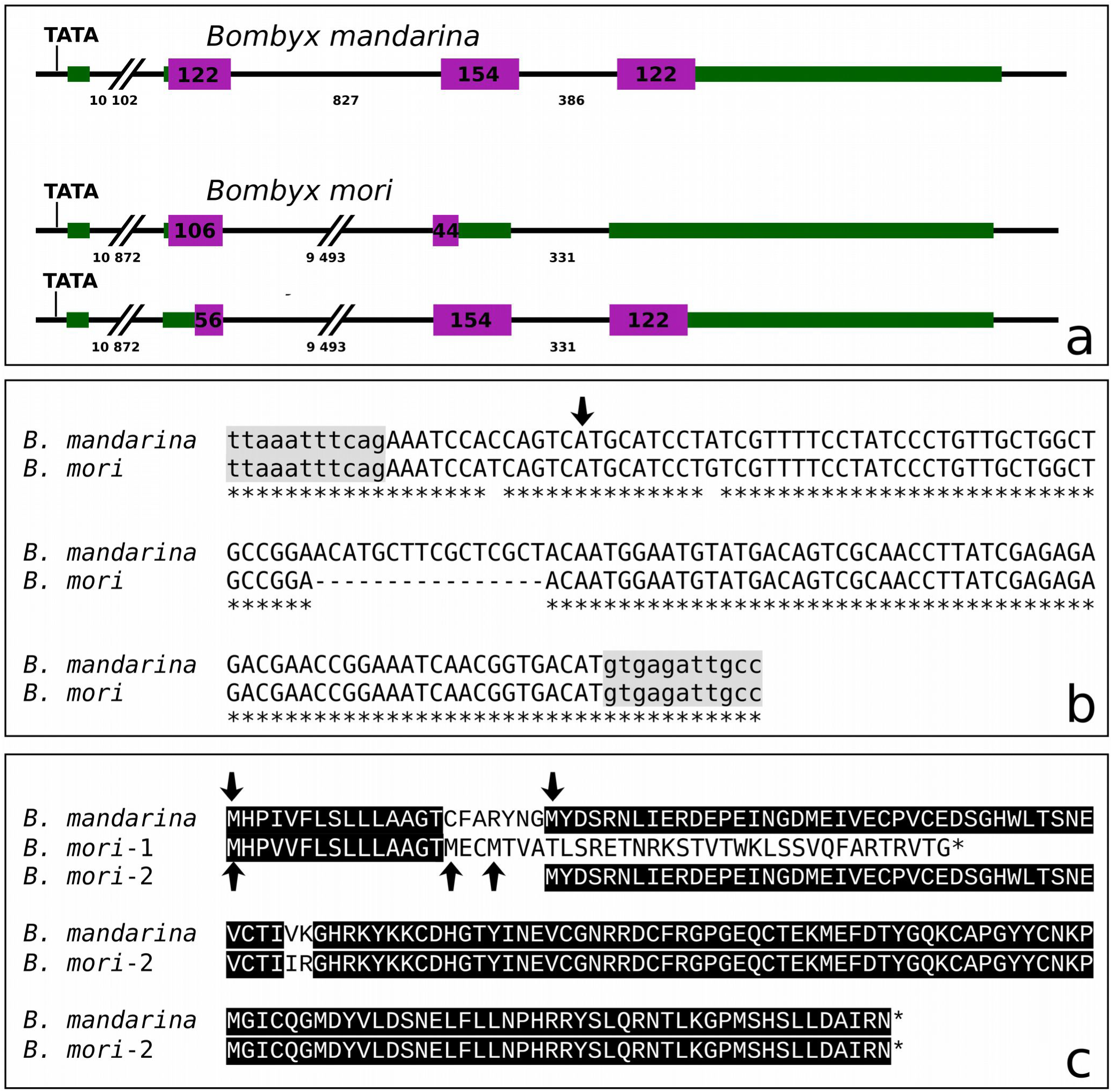
*Bombyx* neuroparsin genes. **a.** Comparative structure of the *Bombyx* neuroparsin genes as present in the genome assemblies. Green and purple indicate exons, green the untranslated and purple the translated parts. The numbers in the purple boxes and below the introns indicate their sizes in base pairs. Both genes have four exons, of which first is untranslated. TATA indicates the location of the TATA box. Note the large difference in the size of the intron between the first and second coding exon. Two different possible translations of the *B. mori* mRNA are represented, that would yield very different translated proteins. **b.** Direct comparison of the second exon (capitals) and the sequences immediately surrounding them (lower case). The start codon is indicated by an arrow. Note that apart from two SNPs, one before the start codon and one that causes a ile-val change, there is a 16 nucleotide deletion in the *B. mori* sequence. **c.** The possible translations of the two genes. In *B. mandarina* a conventional neuroparsin precursor is predicted. In *B. mori*, one predicts a very short peptide without any sequence similarity to neuroparsin but with a signal peptide. In case the first three start codons were to be ignored, a protein could be produced that has a neuroparsin sequence, but lacks that a signal peptide and hence can not be expected to be secreted. Black highlighting indicates identical amino acid residues.

The deletion of 16 base pairs from the first coding exon either leads to a very small peptide with a signal sequence or, in case the first three methionine codons were to be ignored, to a protein that contains the neuroparsin sequence, but lacks a signal peptide (Fig. 4). Therefore this mutation makes it impossible for the neuroendocrine cells expressing this mutated neuroparsin gene to secrete this neurohormone. Over time a large number of different silkworm strains have been sequenced, in order to analyze the domestication of the silkworm and and its underlying evolutionary process (Xia et al., 2009; Xiang et al., 2018). Sequences for still other lines are available at silkbase (http://silkbase.ab.a.u-tokyo.ac.jp). Although the latest *B. mori* genome assembly (Kawamoto et al., 2019) shows a 16 basepair deletion in the neuroparsin coding sequence, only a limited number of silkworm strains have this deletion. Thus in the 161 different *B. mori* strains in which this could be determined, there were only twelve that carried the 16 base pair deletion while four other strains had both alleles. As genome assemblies are all based on single individuals, the actual number of *B. mori* strains where this allele is present is almost certainly larger, but it must be a minority. These numbers are not consistent with the Hardy-Weinberg equilibrium and, hence, in most populations the allele will be fixed.

It seemed interesting to know whether the enormous increase in size of the intron between the first and second coding exons might be of functional significance. I therefore determined, where possible, the size of this intron in other Lepidopteran genomes, as well as the various *Bombyx* strains. Whereas it was relatively easy to determine coding sequences in genome assemblies with large numbers of small contigs, the determination of the size of this intron is more difficult. Nevertheless, there were only a few species in which the size of this intron could not be determined. In the large majority of species, this intron is quite small, and there are only a few species where it is larger than 1500 base pairs; the largest intron was found in *Tuta absoluta* (2285, Table S1). Determining the size of the intron in all the *Bombyx* strains was more difficult, in five *B. mandarina* strains where this could be determined it varied between 827 and 1292. In the thirty *B. mori* strains where this could be established it varied between 918 and 1298; none of these strains had the 16 base pair deletion. There was not a single *B. mori* strain having this deletion for which it was possible to determine the size of the intron, apart from the of the official sequence assembly of the p50T strain (Kawamoto et al., 2019). Nevertheless, in several strains its size had to be at least more than 1900 base pairs and this was found both in strains that have the 16 basepair deletion and strains that do not. Although this suggests that the intron size increased before the deletion occurred, it does not definitively prove it.

## 4. Discussion

*Bombyx mori* has been and still is the model for lepidopteran neuropeptides and it was surprising that neuroparsin that seemed to be produced in large neuroendocrine cells of the brain had escaped detection in this species as well as other Lepidoptera. It is now clear that Lepidoptera do have such a hormone, but its sequence is so different from that of other Arthropod neuroparsins, that it is difficult to recognize as such. Nevertheless, the expression in *Galleria* brain neuroendocrine cells of this protein confirms that it is indeed neuroparsin.

In the *Bombyx* strain used preferentially in the laboratory and the one with the best genome assembly (Kawamoto et al., 2019) the neuroparsin gene shows two mutations. The first one makes it impossible for the putative neuroparsin precursor to have a signal peptide and hence it can not be released as a hormone into the hemolymph. Although, it is also possible that no neuroparsin is made at all, and that only small peptide with a signal peptide is the protein product made from this gene. Either way, this probably explains why neuroparsin has not previously been identified from this species.

It is interesting to note, that there is second mutation in the *Bombyx* neuroparsin gene, which concerns a very significant increase in the size of the intron between the first and second coding exons. In those moths and butterfies where its size could be determined it was never more than 2285, and in most of them it was less than 1000 base pairs. Yet in the sequenced *B. mori* strain it is 9493 base pairs long. It is noticeable that neuropeptide genes that are heavily expressed in small number of neurons, such as the insulins, AKHs, or the *Locusta* vasopressin genes, all seem to have very small introns, suggesting that small intron sizes may facilitate efficient neuropeptide production. It seemed possible that once a 16 base pair deletion effectively inactivated the neuroparsin gene there was no longer any selection for maintaining the small size of this intron. Alternatively, if for some unknown reason the neuroparsin gene has no function in domesticated silkworms, the increase of the size of this intron may have preceded the deletion. Results suggested that the deletion in the coding sequence occurred after the increase of the intron size. If one accepts the notion that small intron sizes facilitate efficient expression of (neuropeptide) genes, it then also suggests that during domestication the importance of the neuroparsin gene was diminished well before the gene was completely inactivated by the deletion in its coding region. Such a scenario may explain the absence of a detectable *in situ* hybridization signal in *B. mori.*

It is indeed surprising, that while in virtually all Lepidopteran genomes a neuroparsin gene can be detected that seems to be functional, it is only in some domesticated *Bombyx* strains that this gene has apparently become superfluous. It is well documented that domestication leads to significant changes in the genome of a species and this also occurred in the silkworm (Xiang et al., 2018). As this mutated gene is only present in some but not all domesticated silkworms it is clearly not a mutation that facilitated domestication, it is rather a gene that domesticated silkworms can do without, while remaining essential in wild moths.

It is obvious that just like *Drosophila melanogaster* the domesticated silkworm survives well without a functional neuroparsin gene. It can thus not be excluded that it is a pure coincidence that this happened in some domesticated strains of *Bombyx mori* and not in other Lepidoptera, assuming that the absence of the gene in *Plutella xylostella* and *Operophtera brumata* is indeed due to technical problems rather than a genuine absence from their genomes.

A recent publication on *Anopheles coluzzii* showed the venus kinase receptor to be required for protection against infection by *Plasmodium* parasites (Gouignard et al., 2019). As the venus kinase receptor can also be activated by L-amino acids (*e.g.* Ahier et al., 2009), this does not necessarily show the implication of neuroparsin. Nevertheless, within in this context it is at least intriguing to note that some of the species with the largest intron sizes all have some type of natural exterior protection during the larval stages which not only makes chemical control of such species difficult, but likely also protects them against viruses, bacteria and perhaps parasites. Thus *Cydia pomonella* (intron size 1598, Table S1) larvae live and feed inside apples, larvae of *Galleria mellonella* (intron size 1884, Table S1) live inside bee hives, *Tuta absoluta* (intron size 2285, Table S1) is a leaf miner and caterpillars of *Megathymus ursus* (intron size 1549, Table S1) feed inside yucca roots and agave leaves (Cong et al., 2019). In any case, a hormone that would be important for protection against parasites or infections, would likely be less important in a domesticated species, as diseased animals would be rapidly eliminated by silk farmers.

The identification of Lepidopteran neuroparsins shows how variable its structure can be in insects and it illustrates the difficulty of finding putative orthologs of this hormone in other Arthropods. Although not limited to insects as the Decapod neuroparsins convincingly show, it remains unknown whether this hormone is present in other Arthropods like *e.g.* Chelicerates. Venus kinase receptors have a wide distribution within Protostomes and even Deuterostomes (Vanderstraete et al., 2013; Dissous et al., 2014). Since neuroparsin can activate this type of receptor (Vogel et al., 2015), *a priori*, one might expect neuroparsin orthologs to have a similar wide distribution. However, these receptors are also activated by amino acids (*e.g.* Ahier et al., 2009; Vanderstraete et al., 2014; Gouignard et al., 2019) and hence the distribution of neuroparsins could be much more limited. It is therefore regrettable that the extreme variability of this hormone in insects suggests, that it will be very difficult to identify neuroparsin orthologs, if they even exist.

## Acknowledgements

I thank Michal Žurovec for *Bombyx mori* pupae and Dušan Žitňan for his *in situ* hybridization protocol.

