## SupplementaryData for "The hormone neuroparsin seems essential in Lepidoptera but not in domesticated silkworms"

**Jan A. Veenstra**

#### Content

| | |
|---|---|
| Table S1. Lepidopteran Neuroparsin precursors | page 2 |
| Table S2. Presence of neuroparsin specific reads in selected Bombyx mori SRAs | page 8 |
| Figure S1. DNA sequence for *Galleria mellonella* neuroparsin cDNA product sense primer | page 9 |
| Figure S2. DNA sequence for *Galleria mellonella* neuroparsin cDNA product anti-sense primer | page 10 |
| Figure S3. DNA sequence for *Bombyx mori* neuroparsin cDNA product sense primer | page 11 |
| Figure S4. DNA sequence for *Bombyx mori* neuroparsin cDNA product anti-sense primer | page 12 |

### Table S1. Lepidopteran Neuroparsin precursors

Predicted neuroparsin sequences from Lepidoptera. Most of these sequences are predicted from the genome assemblies at NCBI with a smaller number identified from transcriptome assemblies. X in sequences indicates gaps in the genome assemblies, - in sequences indicate incomplete transcript sequences making it impossible to know the N- and/or C-terminal sequences of the precursors. Signal peptides predicted by SignalP-5.0 (http://www.cbs.dtu.dk/services/SignalP/) are highlighted in yellow, while cysteine residues are highlighted in orange. Numbers after a species name in indicate the size of the intron between the first and second coding exons. In a few cases their size could not be determined, while in the case of sequences only known from transcripts the GenBank identifiers are shown. Note that neuroparsin precursor sequences could not be found in a few genome assemblies, in some cases these genes could be assembled using SRAs containing genomic sequences of these species, however no neuroparsin sequences could be identified for two species, *i.e. Operophtera brumata* and *Plutella xylostella.*

***Achaea janata*** NCBI: GHGZ01102743.1
-----
GSYDTRNFIAREEPMPDDDGEYVECPVCDESEHWLGSNENCPIKRGNKTYKKCSLGTYVNEVCGKKLDCYRGPGEQCTVKRE
FDMFGRKCAPGFYC

***Adoxophyes honmai*** 324
MHPIIYLALIAASSCMAHYHSFENHLVEREDPQDADSDLVECPVCEESTWLPNNESCNVLRGNKLYKRCPLGVYINNVCGNR
RDCYRGPGEQCTEKMEVDVYGQKCGHGLYCNRALGKCTGSKPSYNFLYKINRRFCDSPYNIYIQIIQYVFCMYVYSIKLSCQ
SPEG

***Adoxophyes orana*** NCBI: GGMW01028324.1
-----
EESTWLPNNESCNVLRGNKLYKRCPLGVYINNVCGNRRDCYRGPGEQCTEKMEVDVYGQKCGHGLYCNRALGKCTGFGFTVD
SNVQYMINPHRYPLRAGGDI

***Agrotis ipsilon*** 1482
MHSLMYLMLVAVSSCLAHYRGSYESRNFITREEPVPVEGVLVECPVCEDSDHWMASKEECNITKGNKSYKICKFGTYINEVC
GKRLDCYRGPREQCTEKREFDMYGKKCAPGYYCHKWAGVCTGPEYRLDSNWQWTLIPFTGRRSELKVQPEDAKFPFEGLR

***Amylois transitella*** 407
MHPVVYALLLVGSCLAHYRGGYESRLNLIEREEPQEEQDGDIVECPVCGGSGNSEWLPSNEVCPIVRGDKRYKRCQLGTYVN
TVCDNRLDCYRGPREQCTEKMDFDIYGQKCAPGYYCNKYLGVCTGLEYNVESKTQWLLNPNHRYPLRARRSI

***Arctia plantaginis*** NCBI: GGLW01067254.1
-----
GPYDNHNLIVREEPYSGDDRDIVECPVCEESENWLPNTEKCPIKRGDKIYKKCKLGTYINEVCGSRLDCYRGPGEQ-----

***Bicyclus anynana*** 412
MHLVLSIVLLISTSCLAHYRGYESHNLIERDELRDSDGSVLVECPVCELSSSGIPSREVCDIPKGNKMFKRCRRGTYVNEVC
GNRLDCYRGPGEQCTEKMDFDYYGQKCARGFHCDDTFHECFGRGYTPDSRQRFFLSHMRSGIYGHTKKNEIPENSPLYK

***Bombyx huttoni*** impossible to determine intron size
MHPIVILSLLLAAGTCFARYNGIYNNRNLIERDEPEFNSDMEIVDCPVCEDSGYWLSSNEVCNVTKNHRKYKQCNHGTYINE
VCGNRRDCFRGPGEQCTEKMEFDTYGQKCAPGYYCNKPMGICQGMDYVLDSNELFLLNPHRRYFLQRNSLKGSMSHSLLDAI
GN

***Bombyx mandarina*** 827
MHPIVFLSLLLAAGTCFARYNGMYDSRNLIERDEPEINGDMEIVECPVCEDSGHWLTSNEVCTIVKGHRKYKKCDHGTYINE
VCGNRRDCFRGPGEQCTEKMEFDTYGQKCAPGYYCNKPMGICQGMDYVLDSNELFLLNPHRRYSLQRNTLKGPMSHSLLDAI
RN

***Calephelis nemesis*** 541
MQLAAFFLLLALSSSLAHYTGYETKNLIVREEPLLEVEGNLVECPVCEPYSLLPSHEECNIVRGNKKYKRCPLGTYINEVCG
DRLDCYRGPGQQCTEKMDFDAYGQKCAHGYYCDGTFHVCIGVDYTINSHLRWLVNHAYRHPLRRTKEQPDISSLLI

***Calephelis virginiensis*** 2026
MQLAAFSLLLALSSSLAHYTGYETKNLIVREEPLLEVEGNLVECPVCEPYSLLPSHEECNIVRGNKKYKRCPLGTYINEVCG
DRLDCYRGPGQQCTEKMDFDAYGQKCAHGYFCDGTFHVCIGVDYTINSHLRWLVNHAYRHPLRRTKEQPDISSLLL

***Calycopis cecrops*** 238
MPSAIKILLLAVATCAAHYPGYDARGLIIREEPIIEEEGILVECPVCNEISSAPRIMEECPIVLGNKKYKRCIRGTYINEIC
GNRWDCYRGPSEQCTEKMNNDVYGQRCAYGYYCDDKDHVCTGSAFSGKFLINPVNRGHRYPLLAFRHEQTDINDK

***Carposina sasakii*** NCBI: GGMY01049356.1
MYGLILTLLACSVCLAHYRGGYESRNIIEREEPLNYNEGITVPCPVCENTDLWLPSSEECNLVKGGKKYKRCKLGTYVNDVC
GDRLDCYRGPGERCTEKMDVDRLGERCAPGYYCNKAMGVCTGMDFIIDSNELLLLNPNRFPLRERRYAY-

***Cecropterus lyciades*** impossible to determine intron size
MFSLVLFLMLVGACSAHYRGYDGRNLIVREEPSGVAGVLVECPVCENTEDLLSNHEKCDITKGGKKYKMCKRGTYVNEVCGN
RLDCFRGPGEQCSEKMRYDVYGQKCARGFYCNDVFHVCTGLGYNNIDSNALWYLNNLYRYPLGSQHDQTSKVESMLLS

***Chilo suppressalis*** 466
MYTVGFLILAVGTCTAYYGGGYEGRNLIAREEPQEDEGGLVECPVCVNARNGEWVPSNEVCPIVKGNKRYKRCQRGIYVNEV
CDNRLDCYRGPREPCTEKMDFDIFGQKCAPGYYCNKFLGVCTGLEYNPASKTQWVLNPHRRYTLREEHSV

***Colias eurytheme*** NCBI: GGJZ01013362.1
MHHLLYILLAASTSCMAHYRGYESRNLIEREEPHPVDGTLVECPVCESSLSWLPNHEECNIVKGEKRYKKCERGTYINEVCG
GRRDCYRGPGEHCTEKMDYDVYGQKCAHGYYCDNNFNVCTGLGYAIDSHVRWILNHPYRYPLRSQNEEDALNSKPLMLLA-

***Cydia pomonella*** 1598
MHPLLYLMLLAGPCLARYRFEPRNLIERDDLQEKGEWVDCPVCETPSYSSLTSDEVCEMHNGRPAKLCLHGTYINKVCGSRR
DCYRGPDEPCTEKLDNDEFGWKCAPGSTCSSVTGKCIGQ

***Danaus chrysippus*** 506
MHLSIYILASVFYLTTAHYRGYESRNLIVREEPFDLEGLVECPVCQYQVSWLPTREVCNITIGNKKYKRCRMGNYTNTVCGN
RLDCFRGPGEQCTESMENDYYGQKCARGYDCDGTFHVCTGYGYTINSHMRWLLNHVSRYPGNQNDDQLREKALYLA

***Danaus plexippus*** 505
MHLSIYILASVFYLTTGHYRGYESRNLIVREEPFDLEGLVECPVCQYQVSWLPTREVCNITIGNKKYKRCRMGNYTNTVCGN
RLDCFRGPGEQCTESMENDYYGQKCARGYDCDGTFHVCTGYGYTINSHMRWLLNHVSRYPGNQNDEQLREKALYLA

***Eogystia hippophaecolus*** NCBI: GFBH01072795.1
MHPLVYLMLFAGSACLAHYRDGYDSRNLIERDEPQNDVEGELVECPVCENSGNWMSNGEVCPIARGNKRYKKCELGTYVNEV
CGNRLDCYRGAGDRCTEKMEVDPYGQKCAHGYYCNKVLGVCTGLNYAVDSNVQWLINPHRRYPLRANRDI

***Eumeta japonica*** 760
MQLLRLMVICAAVGYSAAYYHDSPQGLGLRDVIEPRDVIENYETAVQKRDEYVLQLELVACPVCDQFLDDECTVLKNGKRYK
KCAFGTYINEVCGNQRDCYRGPKQYCTEKMAEDKLGAKCAHGYTCNAQLHECVGLFNELDSLQLAGLRHYYILPEDNTNSYI
DKYRLARSHHLY

***Galleria mellonella*** 1884
MHPVLYVMLTIGVCLAHYRGGYDARNLIEREEPQDDIEGEIVECPVCIGSSSNDWLPSNELCPIVRGNKRYKRCQRGTYVNT
VCDNRLDCYRGPKEQCTEKMDFDIYGQKCAPGYYCNKYLGVCTGLEYNVESKTQWLLNPNRRYPLRAKRDV

***Heliconius cydno*** 220
MHLVFCLLFLVGSSAAHYHGYESRNLIEREEPNTDMDSLVECPVCETSQDHKECNIDRGNKRFKRCHWDTYFNEKCGNRLDC
YRGPGETCTEKMENDIYGQKCAHGCYCDIAVHQCVCVGYKLDQYLRLALDNPLYRYPIPKSQFEKSVFLENLPKPVMVLA

***Heliconius elevatus*** 225
MHLVFCLLFLVGSSAAHYHGYESRNLIEREEPNTDMDSLVECPVCETSQDHKECNIDRGNKRFKRCHWDTYFNEKCGNRLDC
YRGPGETCTEKMENDIYGQKCAHGCYCDIAVHQCVCVGYKLDQYLRLALDNPLYRYPIPKSQFEKSVFLENLPKPVMVLA

***Heliconius ethilla*** 219
MHLVFCLLFLVGTSVAHYHGYESRNLIEREEPNTDMDSLVECPVCETSQDHKECNIDRGNKRFKRCHWDTYFNEKCGNRLDC
YRGPGETCTEKMENDIYGQKCAHGCYCDIAVHQCVCVGYKLDQYLRLALDNPLYRYPIPKSQFEKSVFLENLPKPVMVLA

***Heliconius hecale*** 224 not found in genome assembly, assembled from SRR7162651
MHLVFCLLFLVGSSAAHYNGYESRNLIEREEPNTDMDSLVECPVCETSQDHKECNIDRGNKRFKRCHWDTYFNEKCGNRLDC
YRGPGETCTEKMENDIYGQKCAHGCYCDIAVHQCVCVGYKLDQYLRLALDNPLYRYPIPKSQFEKSVFLENLPKPVMVLA

***Heliconius hecuba*** 204
MHLVICLLLLAGSSAAHYRGFESRNLIEREEPNTDMDSLVECPVCEVSQDHKECNIDRGNKRFKRCHWDTYFNEKCGNRLDC
YRGPGETCTEKMENDIYGQKCAHGCYCDSAVHQCVCVGYKLDQYLRLVLDNPLYRYPIPKSQFEKSVFLENLPKPVMVLA

***Heliconius heurippa*** 219
MHLVFCLLFLVGSSAAHYHGYESRNLIEREEPNTDMDSLVECPVCETSQDHKECNIDRGNKRFKRCHWDTYFNEKCGNRLDC
YRGPGETCTEKMENDIYGQKCAHGCYCDIAVHQCVCVGYKLDQYLRLALDNPLYRYPIPKSQFEKSVFLENLPKPVMVLA

***Heliconius hierax*** 249
MHLVICLLLLAGSSAAHYRGFESRNLIEREEPNTDMDSLVECPVCETSQDHKECNIDRGNKRFKRCHWDTYFNEKCGNRLDC
YRGPGETCTEKMENDIYGQKCAHGCYCDSAVHQCVCVGYKLDQYLRLALDNPLYRYPIPKSQLEKSLFLENLPKPVMVLA

***Heliconius ismenius*** 226
MHLVFCLLFLVGSSAAHYHGYESRNLIEREEPNTDLDSLVECPVCETSQDHKECNIDRGNKRFKRCHWDTYFNEKCGNRLDC
YRGPGETCTEKMENDIYGQKCAHGCYCDIAVHQCVCVGYKLDQYLRLALDNPLYRYPIPKSQFEKSVFLENLPKPVMVLA

***Heliconius melpomene*** 219
MHLVFCLLFLVGSSAAHYHGYESRNLIEREEPNTDMDSLVECPVCETSQDHKECNIDRGNKRFKRCHWDTYFNEKCGNRLDC
YRGPGETCTEKMENDIYGQKCAHGCYCDIAVHQCVCVGYKLDQYLRLALDNPLYRYPIPKSQFEKSVFLENLPKPVMVLA

***Heliconius numata*** 230
MHLVFCLLFLVGSSAAHYHGYESRNLIEREEPNTDMDSLVECPVCETSQDHKECNIDRGNKRFKRCHWDTYFNEKCGNRLDC
YRGPGETCTEKMENDIYGQKCAHGCYCDIAVHQCVCVGYKLDQYLRLALDNPLYRYPIPKSQFEKSVFLENLPKPVMVLA

***Heliconius pachinus*** 220
MHLVFCLLFLVGSSAAHYHGYESRNLIEREEPNTDMDSLVECPVCETSQDHKECNIDRGNKRFKRCHWDTYFNEKCGNXXXX
XXGPGETCTEKMENDIYGQKCAHGCYCDIAVHQCVCVGYKLDQYLRLALDNPLYRYPIPKSQFEKSVFLENLPKPVMVLA

***Heliconius timareta*** unknown intron size, but likely very small gap in contig
MHLVFCLLFLVGSSAAHYHGYESRNLIEREEPNTDMDSLVECPVCETSQDHKECNIDRGNKRFKRCHWDTYFNEKCGNRLDC
YRGPGETCTEKMENDIYGQKCAHGCYCDIAVHQCVCVGYKLDQYLRLALDNPLYRYPIPKSQFEKSVFLENLPKPVMVLA

***Heliconius wallacei*** 248
MHLVFCLLFLVGSSAAHYHGYESRNLIEREEPNTDMDSLVECPVCETSQDHKECNIDIGNKRFKRCHWDTYSNEKCGNRLDC
YRGPGETCTEKMENDIYGQKCAHGCYCDSAAHQCVCVGYKLDQYLRLALDNPLYRYPIPKSQYEKSVFLENLPKPVMVLAS

***Heliconius xanthocles*** 261
MHLVICLLLLAGSSAAHYRGFESRNLIEREEPNTDMDSLVECPVCETSQDHKECNIDRGNKRFKRCHWDTYFNEKCGNRLDC
YRGPGETCTEKMENDIYGQKCAHGCYCDSAVHQCVCVGYKLDQYLRLALDNPLYRYPIPKSQFEKSLFLXXXXXXXXXXXX

***Helicoverpa armigera*** 492
MHTLVFLMLVTYSSCLAHYRGSYESRNYITREEPKGVLIECPVCPSEGWDAIEEVCIKEGDKSYKLCRFGTYINTACNNRLD
CLRGPREKCTEKHVFNRSNKKCTFGYECNQKEGVCTGPENRPEIYWPYRFGLYPDRRSSSKSTAEEAFEDMTN

***Helicoverpa zea*** 474
MHTLVFLMLVTYSSCLAHYRGSYESRNYITREEPKGVLIECPVCPSEGWDAIEEVCIKEGDKSYKLCRFGTYINTACNNRLD
CLRGPREKCTEKHVFNRSNKKCTFGYECNQKEGVCTGPENRPEIYWPYRFGLYPDRRSSSKGTAEEAFEDMTN

***Heliothis virescens*** 469 NOT IN GENOME assembled from SRR5463746
MHTLVFLMLITFSSCLAHYRGSYESRNFVTREEPTEVLVECPVCPSEGWDAIEEVCIKEGDKSYKLCKFGTYVNTACNNRLD
CLLGPREKCTEKHVFNRSNKKCTYGYVCNEKEGVCTGPEFHVSNTWRYFVEPYSGLRSSKERPEAAFEEMRN

***Hyphantria cunea*** 712
MHPLLYLMLLTGFCLAHYRGSYDNHNLIAREEPVAEDDGDIVECPVCENSEHWSPNTEKCPIRRGNKVYKKCNLGTYINEVC
GSRMDCYRGPGEQCTVKREFDNYGRKCAPGYYCHKSLGVCIGLGYQIDSNVQWQLNRFPYTWRRSDLKGISDESKFSFEGGP
N

***Hyposmocoma kahamanoa*** 238
MYLFLYTMLLTGGGALAHFRGYTTRNLFVRDEPVNQVEEINFVECPVCNNTSLWPPLIENCEVNLNGKRYKQCKFGTYKNRV
CGNRNDCKRGPGDKCSTFDNRDMYDLRCAPGYHCDGLEHRCIGGSNNPNPVNRYYLSPSSFSGSDYTRSTDLYNSREKRSAP
YSFFAPH

***Laparus doris*** unknown intron size, but likely very small gap in contig
MHLVICLLLLAGSSAAHYRGFESRNLIEREEPNTDMDSLVECPVCETSQDHKECSIDRGNKRFKRCHWDTYFNEKCGNRLDC
YRGPGETCTEKMENDIYGQKCAHGCYCDSAVHQCVCVGYKLDQYLRLALDNPLYRYPIPKSQFEKSLFLENLPKPVMVLA

***Leptidea sinapis*** 1170
MQPILCILLALLASSTAHYRGYEDRNLIERDEPNISDGTLVECPVCENTSWLPSREQCNIVKGNKRYKKCERGTYVNEVCGN
RLDCYRGPNEQCTEKMDFDVYGQKCGHGYYCDRTLHVCTGLGYTIDSKIRLLLNDLQGFRYPLHPNLDEDSPQKSMLLLA

***Lerema accius*** 1107
MNFLIVITILAGACVAHYRGGYDNRNLIEREEPDMPGVLVECPVCENEDGLLPINEKCNIIKGGKRFKKCERGTYVNEVCGN
RIDCYRGPGEKCSEKMAFDDYGIKCARGFYCNDVFHVCTGPGFTIDSNSLWLLNGVYRYPLPFRQQERAAKSESAMILA

***Loxostege sticticalis*** NCBI: GFCJ01054349.1
-----
QDDMGGELVECPVCLGARNGEWLPSNEMCPILRGNKRYKRCQRGTYINEVCDNRLDCYRGPREQCTEKMDFDIYGQKCAPGY
YCNKYLGVCTGLEYNPDSKTQ

***Lymantria dispar*** 992
MHSLLYLLLLTGFCLAHYRESYDSRNFIVREEPSNSFDGEIIECPVCEDPESWMPGEKCPIVKDNKTFMKCKYGTYINEVCG
GRLDCYRGPGEQCTVKWAFDPYGKKCAPGYICNKSLGICSGYTYHIDSNIQRQLNHFPYTFRRSDLKDLSEEPKLNFEIAQI

***Mamestra configurata*** 729
MHSLVFLMLIALSSCLAHYRGSYDSHNFITREEPTAVDGVLVECPVCEQSDHWMASNVEECPIRKGNKTYKICKFGTYINEV
CGKRLDCYRGPLEQCTEKRVFDMYGKKCARGYYCNQLAGVCTGPEVGDRLDSIRQWQLIPFTGRRSELKEQSEDTKFFEGPR
N

***Manduca sexta*** from transcriptome, genome assembly problem
MQSLVYLMLIIAGGVLAHYRGVYDSRNLIEREERMFDGDVEIVECPVCVDSGHWLPSNEECTIRKGNKKYKKCELGTYINEV
CGNRRDCFRGAEEQCTEKMEVDIYGQKCGAGYYCNKLMGICQGLEYVPDSTQQFLLNPHHRYSLQRSGIKTRAANHFLDTIR
N

***Megathymus ursus*** 1549
MHSLIIIMILVGAFVTNAHYRGEYENRNLIEREEPDAPGMLVECPVCENMDGLLPINEKCNIIKGGKRYKKCERGTYVNEVC
GNRIDCYRGPGEKCSEKMTFDDYGIKCARGFHCHDVLHVCTGLGYTIDSNTLWVLNGVYRYPLSFRSQDRPPTKSESALILA

***Meliteaea cinxia*** 386
MHSVIYILLIVGNCLAHYRDYESRNLIERDEPQLDMEGIVECPVCENTHRWLLPQEKCDIIIGNKRFKKCTMGTYNNEKCGN
RLDCFRGPGETCTEKMEDDIYGQKCARGYYCNGGLHVCTGLGYTVDPFFILTSRYHRYPYQNKTYLKDILEKSALLFA

***Mythimna separata*** NBI: GFCT01035668.1
MHSIVFLMLVALSSCLALYRGSYESRNYITREEPVAVDGYKVLVECPVCEHTDHWMASNVEECPIRKGNKTYKICKFGTYIN
EVCGKRLDCFRGPLEQCTEKREFDMYGKKCAPGYYCNKLAGVCTGPDVSDRLDSIRQWQLIPFTGRRSELKEDDKFSFDGLR
N

***Neruda aoeda*** incomplete, poor genome assembly
MDLVICLLLLVGSSAAHYHGYQSRNLIEREEPNTDMDSLVECPVCETSQDHKECNIDRGNKRFKRCHWDTYFNEKCGNRLDC
YRGPGETCTEKMENDIYGQKCAHGCYCDKAVHQCXXXXXX

***Ostrinia furnacalis*** 354
MHTLVFFVLATSTCLAYYGGYDSRNLIVREEPQDEMGGELVECPVCLGARNGEWLPSNEVCPIMRGNKRYKRCPRGTYVNEV
CDNRLDCYRGPREQCTEKMDFDVYGQKCAPGYYCNKYLGVCTGLEYNVDSKTQWLLNPNRRYPLRARRAA

***Ostrinia nubilalis*** NCBI: GAVD01034356.1
-----RLDCYRGPREQCTEKMDFDVYGQKCAPGYYCNKYLGVCTGLEYNVDSKTQWLLNPNRRYPLRARRAA

***Papaipema sp.*** NCBI: GGEH01023339.1
MRSLMYLMLFAFSSCLAHYRGSYESRNFITRDEPAPVEGVLVECPVCEELSSEHWTASNEECPVRQGNKTFKICKLGTYINE
VCGKRLDCYRGPREQCTEKREFDLYGQKCAPGYICNKWAGVCTGPDFRLGNNWPWPLVPFTGRRSEFKIQPEDAKFSFEEIR
K

***Papilio glaucus*** 212
MHLLAYILLIVSASCLAHYRVGYENHRMIQRDEPHDDLEGVLVECPVCENNGLLPSHENCNIIRGNKKFKKCEFGTYINEVC
GNRHDCYRGPAEQCTEKMEVDIYGQKCAHGYYCDNYFNVCVGLGYTVDRHVQWIMNHLYQYPLRDDKDISKPKSSVMLIV

***Papilio machaon*** 261
MHLLAYILFFAATSCCLAHYRGGYESHRMFERNELHEDVDGVLVECPVCESNGVLPTHEHCNIIRGNKKYKKCEFGTYKNEV
CGNRRDCYRGPAEQCTEKMEIDVYGQKCAYGYYCDNLYNVCVGLDYSVERPFGIYNHLYQYPLRDDKLIV

***Papilio memnon*** 205
MHLLAYILLIAAASCMAHYRGGYESHRMFERDELHEDLDGVLVECPVCESNGLLPTHEQCNIIRGNKKYKKCEFGTYKNEVC
GNRRDCYRGPAEQCTEKMEIDVYGQKCAYGYYCDNLYNVCVGLDYSVERPLGVYNHLYQYPLRDDKLIV

***Papilio polytes*** 217
MHLLAYILFLTATSCMAHYRGGYESHRMFERDELHEDLDGILVECPVCESNGLLPTHEHCNIIRGNKKYKKCEFGTYKNEVC
GNRRDCYRGPAEQCTEKMEIDAYGQKCAYGYYCDNLYNVCVGLDYSVERPLGVYNHLYQYPLRDDKLIV

***Papilio polytes*** NCBI: GGKB01058304.1 one amino acid difference with genome
MHLLAYILFLTATSCMAHYRGGYESHRMFERDELHEDLDGILVECPVCESNGLLPTHEHCNIIRGNKKYKKCEFGTYKNEVC
GNRRDCYRGPAEQCTEKMEIDVYGQKCAYGYYCDNLYNVCVGLDYSVERPLGVYNHLYQYPLRDDKLIV

***Papilio xuthus*** 369
MHLLAYILFFAATSCCLAHYRGGYESHRMFERDELHEDVDGVLVECPVCESNGVLPTHEHCNIIRGNKKYKKCEFGTYKNEV
CGNRRDCYRGPAEQCTEKMQVDVYGQKCAYGYYCDNLYNVCVGLDYSVERPFGIYNHLYQYPLRDDK

***Pararge aegeria*** 1065
MHQLLAIILLVSTTCLAHYRSYDSHNLIERDEPRLSDGILVECPVCEQSPSGIPTREVCDIPKGNKMYKKCRRGTYINEICG
NRLDCYRGPGEQCTEKMDYDYYGQKCARTFNCDGLFHECIGLGYTPDSRQRFFLSQMHSGIYGQAKKNDLRENSLMLYA

***Phoebis sennae*** 536 not found in genome assembly, assembled from SRR3091617
MHPLLYILLAASTCMAHYRGYEDRNLIEREEPHIDGTLVECPVCDSSPSWLPSHEECNIIKGEKRYKKCERGTYINEICGGR
RDCYRGPGEHCTEKMNYDIYGQKCAHGYYCDNTFHVCTGLDYKIDSHTRWFFNHDNRYPLRSQNDEDSLNAKSLMLLA

***Phyllodes eyndhovii*** NCBI: GCOO01011652.1
-----
NNERCNLKRGDKTYKKCRLGTYINEVCGNRLDCYSGPGEQCTVKRVFDTYGKKCAPGYYCNPTLGLCTGLEYHANSNLQSQL
NHYPYEAKFNYEGNRN

***Pieris rapae*** 333
MSPILYLLLVSAGCMAHYRGYQGRNLIEKEEHHPNEGILVECPVCYNAPWQFNPDNGNPKCNITIGAQRYKKCDRGSYFNEV
CGRRDCYRGPGEPCTEKMEEDEYGAKCAFGYDCDFSYHVCTGIGFSISIHDRLSFPRLYPLRPLNEGENQAKSMLLT

***Plodia interpunctella*** 785
MHPAMCALLLAGCCLAHYHGGYEARSNLIEREEPQDDLDGDLVECPVCVGSSNSEWLPSNEVCPIVRGDKRYKRCQLGTYVN
TVCDNRLDCYRGPKEQCTEKMDFDIYGQKCAPGYYCNKYLGVCTGLEYNVESKTQWLLNPHHRYPLRARRSI

***Polyommatus icarus*** NCBI: GAST02007939.1
MHMVIQLLVLACACAAHYTGFDSRDLIIREEPADEDGTLVECPVCENVLSFLPSHEECPIIRGNKKYKKCLKGTYINEVCGN
RLDCYRGPREQCTENMNFDAYGQKCAHGYYCDNTFHVCTGSDFAIESKYHWLIHPGYRFPLNTINHDRPEIKSIILA-

***Spodoptera frugiperda*** 573
MQSLVYLMLVAFSSCQAHYRGSYDSRNFITREEPTPKGVLIECPVCDQTDHWMASTEDCPIKNGSKAFKICRFGTYINEVCG
ERVDCFSGPGEQCTEKREFDMYGKKCAPGYYCNKMEGVCTGLEFQLGNNWQWKMYPLRSELRNQPDDSKFSFEGVRN

***Spodoptera litura*** 671
MQSLVYLMLVAFSSCQAHYRGSYDSRNFITREEPTTKGVLIECPVCDQTNHWMASTEDCPIRNGSKTFKICKFGTYINEVCG
KRVDCYSGPGEQCTEKREFDMYGKKCAPGYYCNKMEGVCTGLEFQLGNNWQWKMYPYTGVRSELRNQPEDSKFSFEGVRN

***Thaumetopoea pityocampa*** NCBI: GBZB01005364.1
MQALLWLTLVTCGTCLAHYRDTHDSRNFIVREEPLIDDEGEIVECPVCEETEHWMVNHEKCTTVRGNKLYKRCTLGTYRNEV
CGNRLDCYSGPGEVCTDKRELDMYGKKCAPGYYCNKKLGVCTGLGYTVDSNVQWKLNGYPYTWRRDPAEHSDSFAPIRN

***Tischeria quercitella*** NCBI: GENO01134828.1
MWTVLAMVLIAGACSGHYGPSPYKLPFSLTIPEKLVEPKLVECPVCRLESSIECEPFDSVEGRKMLKKCRHGTFPNTMCNNK
TDCYRGPGESCNEDEGIHCVPSLWCNNYVCKGTSDTGAATKTHSLDTFLYPVTWRNGFQ

***Trichoplusia ni*** 200
MHSLVLFMLVTLATCLAHYRDTSDSRNYVLREEPSNNKVVLVECPTCVSATPWLDKMERCPIKKGNKSYKICMHGTYENEQC
GHRLDCYRDANEECTEKREFDNHDKRCRTGYMCNKELGICVGPEGRPDTWQLKLMTIIGRPNDHKTHTVDDGIPFEGL

***Tuta absoluta*** 2285
MWVFVYALLVTGALGHYQLYDPRSIQNDLEPRQEPFGAGQGKFQVVKCPVCESSNSNLGVESCSPGDSKICRYGVYRNPGCG
RPDCFRGEGELCAENPNAEGFVARCAPGYDCQFNRCVNPLVTQHYNVDRIHEQLYRLKQSPVLWNDN

***Vanessa tameamea*** 590

MHPVFFILILAGTCLAHYRGYESRNLIERDEAHAEMEGIVECPVCENSHSWLLPQEECDIVKGNKRFKKCHMGTYLNEKCGN
RLDCYRGPGESCTEKMEYDIYGQKCARGYYCNSALHVCTGLRYTVDPLSHLILSNRLYRYPLHSQNDDLIDKSAFFVA

#### Table S2. Presence of neuroparsin specific reads in selected *Bombyx mori* SRAs

| Number of reads | SRA identifier | Tissue |
|---|---|---|
| 2 | SRR8493716 | midgut |
| 9 | SRR8493715 | fatbody |
| 4 | SRR8493709 | hemocyte |
| 98 | ERR2018562 | male head |
| 115 | ERR2018561 | male head |
| 720 | ERR2018560 | female head |
| 162 | ERR2018559 | female head |
| 5 | ERR2018557 | ovary |
| 4 | ERR2018558 | ovary |
| 4 | DRR095116 | silk gland |
| 17 | DRR095113 | malpighian tubule |
| 6 | DRR068895 | testis |
| 5 | SRR7812746 | integument |
| 25 | SRR7054573 | colleterial gland |
| 3 | SRR6984041 | female antennae |
| 0 | SRR6984042 | male antennae |
| 0 | SRR6984043 | larval mouthparts |
| 2 | SRR6984044 | larval mouthparts |
| 0 | SRR6129052 | ovary |
| 16 | SRR5119600 | prothoracic gland |
| 429 | SRR4425250 | male brain |
| 380 | SRR4425251 | female brain |
| 1219 | DRR023191 | infected brain larvae |
| 900 | DRR023192 | infected brain larvae |
| 946 | DRR023193 | wild type brain larvae |
| 1445 | DRR023194 | wild type brain larvae |
| 226 | SRR1928169 | brain of silkworm |
| 23 | DRR019433 | Brain from wandering stage |

**File: 37HF36_96560334_96560334.ab1**

Sample Name: *Galleria mellonella*
Mobility: KB_3730_POP7_BDTv3.mob
Spacing: 13.839
Comment: sense primer

Signal Strengths: A = 4866, C = 5756, G = 3617, T = 5096
Lane/Cap#: 30
Matrix: n/a
Direction: Native

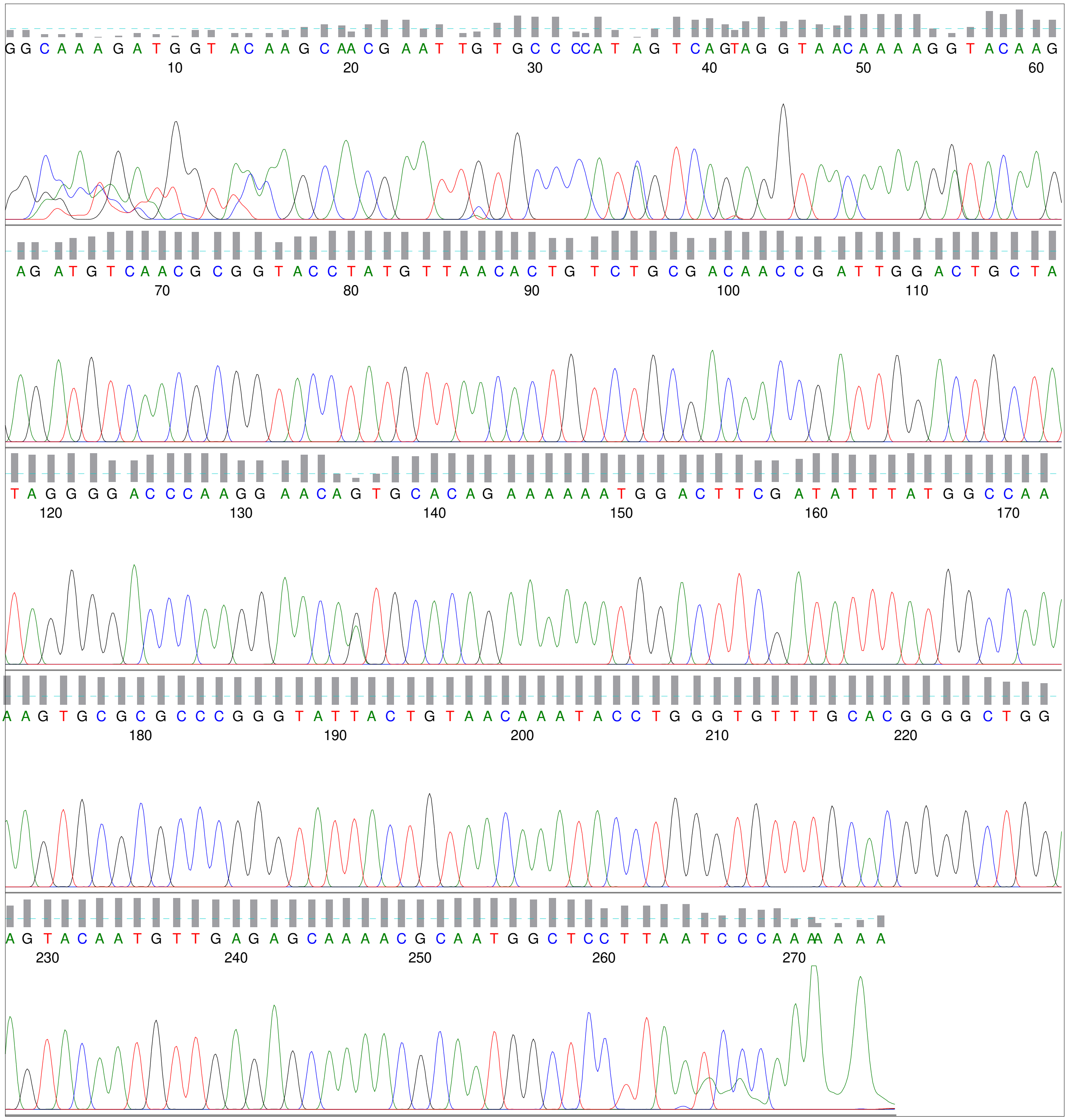

**File: 37HF37_96560532_96560532.ab1**

Geospiza
www.geospiza.com

| | | | |
|---|---|---|---|
| Sample Name: | *Galleria mellonella* | Signal Strengths: | A = 2009, C = 3744, G = 2224, T = 3442 |
| Mobility: | KB_3730_POP7_BDTv3.mob | Lane/Cap#: | 68 |
| Spacing: | 14.4341 | Matrix: | n/a |
| Comment: | **anti sense primer** | Direction: | Native |

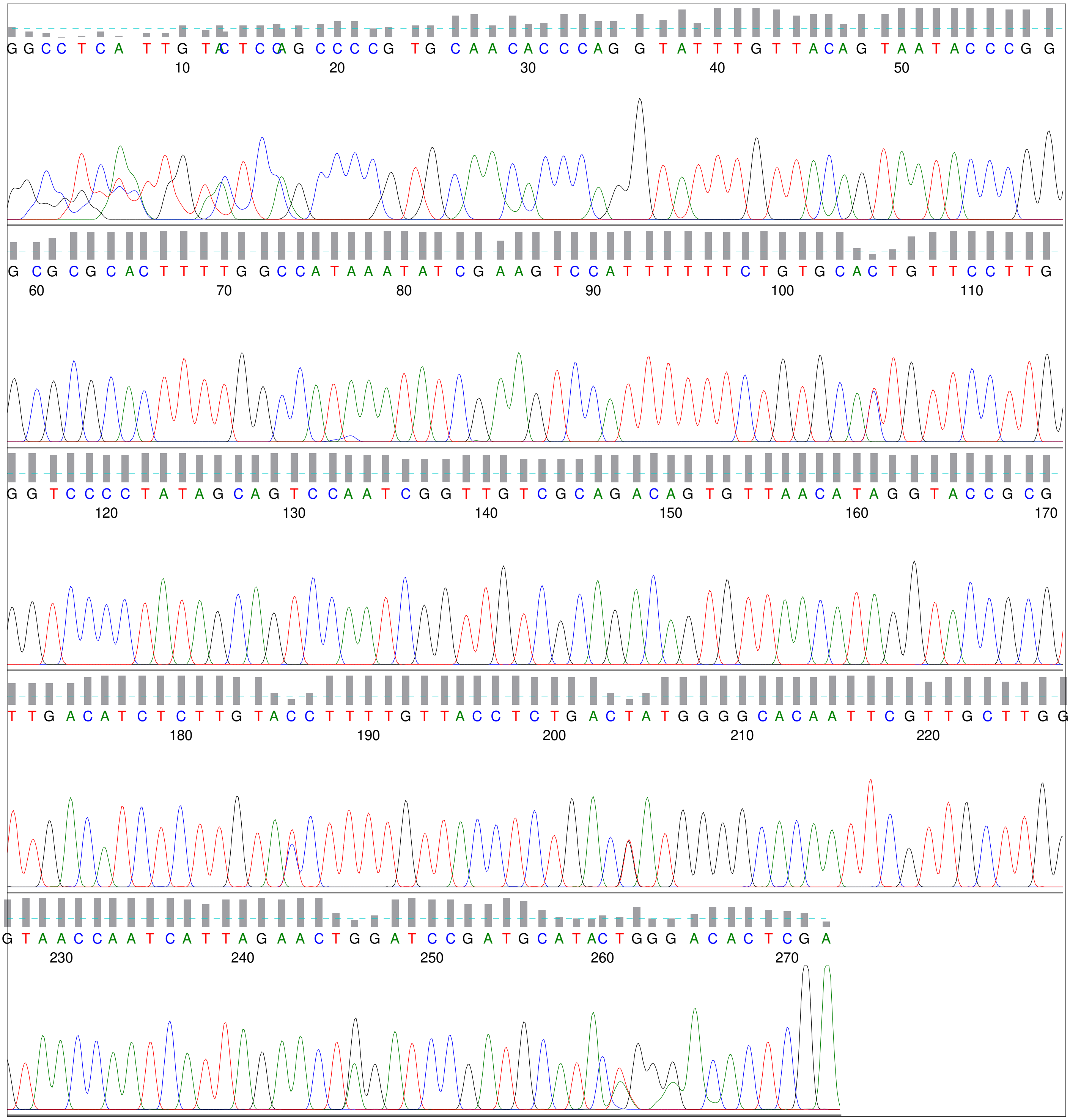

**File: 37HF28_96558324_96558324.ab1**

Sample Name: *Bombyx mori*
Mobility: KB_3730_POP7_BDTv3.mob
Spacing: 13.7387
Comment: sense primer

Signal Strengths: A = 4715, C = 5331, G = 3315, T = 4483
Lane/Cap#: 79
Matrix: n/a
Direction: Native

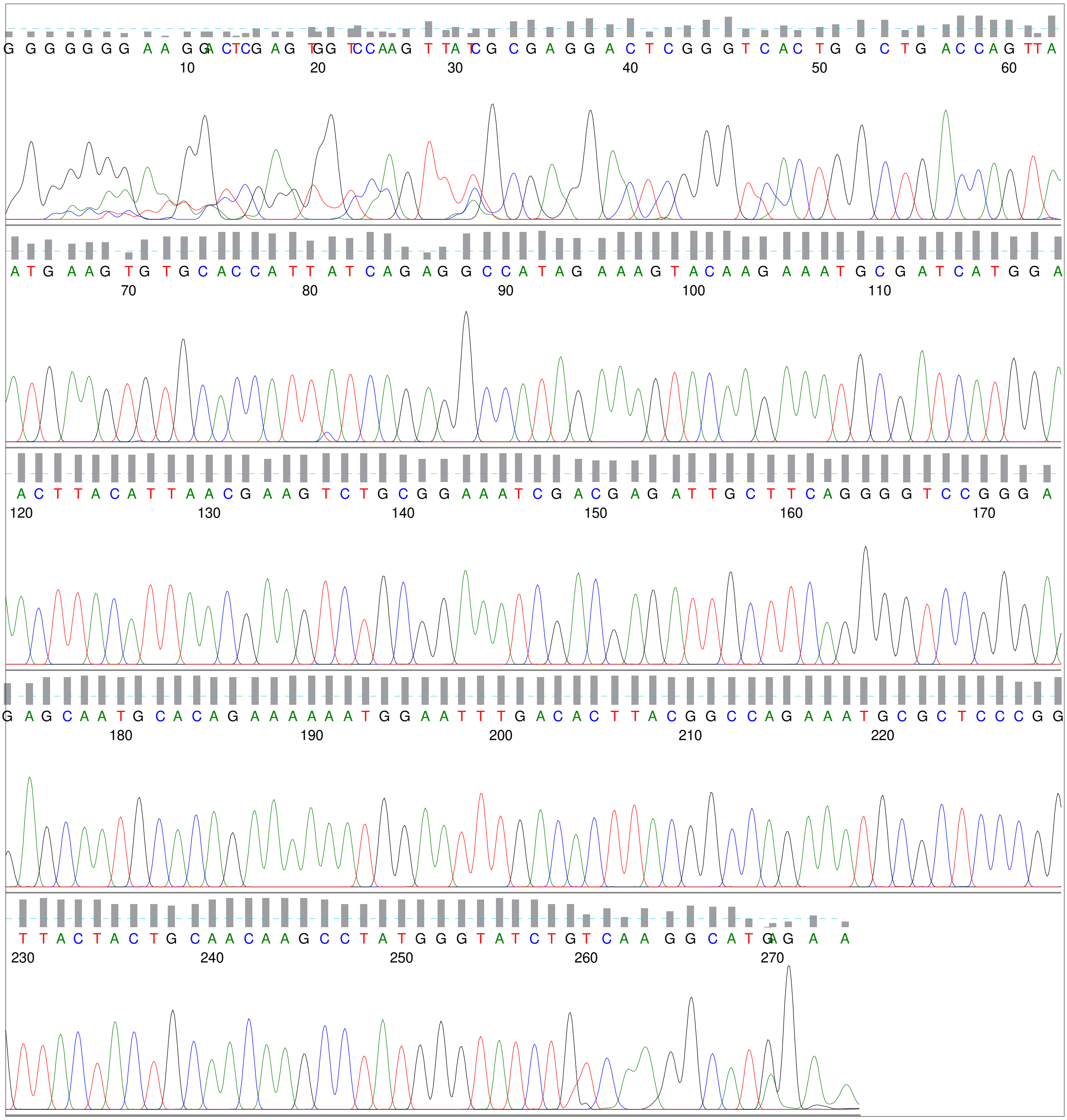

**File: 37HF29_96558874_96558874.ab1**

Sample Name: *Bombyx mori*
Mobility: KB_3730_POP7_BDTv3.mob
Spacing: 13.6917
Comment: anti sense primer

Signal Strengths: A = 4216, C = 6984, G = 3383, T = 6465
Lane/Cap#: 75
Matrix: n/a
Direction: Native

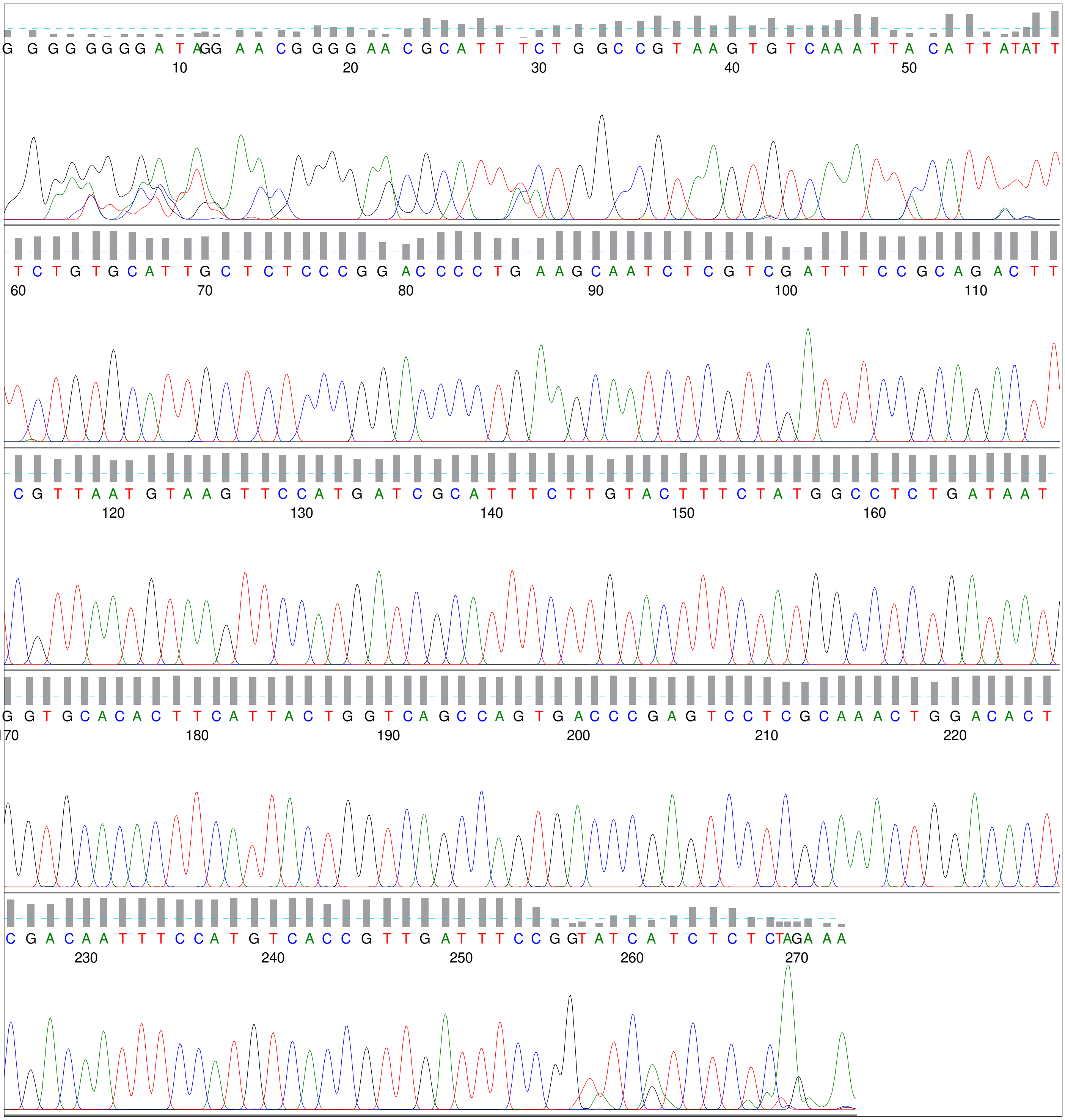
